# The Ecological Forecast Horizon, and examples of its uses and determinants

**DOI:** 10.1101/013441

**Authors:** Owen L. Petchey, Mikael Pontarp, Thomas M. Massie, Sonia Kéfi, Arpat Ozgul, Maja Weilenmann, Gian Marco Palamara, Florian Altermatt, Blake Matthews, Jonathan M. Levine, Dylan Z. Childs, Brian J. McGill, Michael E. Schaepman, Bernhard Schmid, Piet Spaak, Andrew P. Beckerman, Frank Pennekamp, Ian S. Pearse

## Abstract

Forecasts of ecological dynamics in changing environments are increasingly important, and are available for a plethora of variables, such as species abundance and distribution, community structure, and ecosystem processes. There is, however, a general absence of knowledge about how far into the future, or other dimensions (space, temperature, phylogenetic distance), useful ecological forecasts can be made, and about how features of ecological systems relate to these distances. The ecological forecast horizon is the dimensional distance for which useful forecasts can be made. Five case studies illustrate the influence of various sources of uncertainty (e.g. parameter uncertainty, environmental , and demographic stochasticity, evolution), level of ecological organisation (e.g. population or community), organismal properties (e.g. body size or number of trophic links) on temporal, spatial, and phylogenetic forecast horizons. Insights from these case studies demonstrate that the ecological forecast horizon is a flexible and powerful tool for researching and communicating ecological predictability. It also has potential for motivating and guiding agenda setting for ecological forecasting research and development.

## 2 Introduction

Forecasts are statements about what the future may hold in store (Coreau *et al.* 2009), and are useful for all kinds of decision-making, including in economic, political, and personal spheres. Ecological examples include forecasts of species distributions (e.g. Guisan & Thuiller 2005), functional diversity (e.g. Kooistra *et al.* 2008; Schimel *et al.* 2013), phenology (e.g., Cannell & Smith 1983; Diez *et al.* 2012), population size (e.g. Ward *et al.* 2014), species invasions (e.g. Levine & Antonio 2003), agricultural yield (e.g. Cane *et al.* 1994), pollinator performance (e.g. Corbet *et al.* 1995), extinction risk (e.g. Gotelli & Ellison 2006a), fishery dynamics (e.g. Hare *et al.* 2010; Travis *et al.* 2014), water quality (e.g. Komatsu *et al.* 2007), forest carbon dynamics (e.g. Gao *et al.* 2011), ecosystem services (e.g. Homolová *et al.* 2013), disease dynamics (e.g. Ollerenshaw & Smith 1969), and interspecific interactions (e.g. Pearse & Altermatt 2013).

Although forecasting has been part of ecology for decades, current and expected environmental change is motivating ever increasing interest in ecological forecasting. There is a pressing need to deliver information about the probable future state of populations, communities, and ecosystems in order to better inform conservation, management, and adaptation strategies (Clark *et al.* 2001; Sutherland *et al.* 2006; Tallis & Kareiva 2006; Evans 2012; Mouquet *et al.* 2012; Purves *et al.* 2013). Furthermore, accurate forecasting (i.e. correct prediction) is sometimes regarded as the hallmark of successful science (Evans *et al.* 2012), and as such can be a powerful driver of advances in knowledge about how ecological systems work (Coreau *et al.* 2009). This study rests on the premises that accurate ecological forecasts are valuable, that our knowledge about ecological forecasting is relatively sparse, contradictory, and disconnected, and that research into ecological predictability is worthwhile (contrary to, for example, Schindler & Hilborn 2015). Ecologists need to know what properties and components of ecological systems are forecastable, and need to quantify the uncertainties associated with these forecasts (Clark *et al.* 2001; Godfray & May 2014). A systematic understanding of forecast performance in relation to modelling practices, sources of uncertainty, organismal characteristics, and community structure can guide ecology to become an even more predictive science.

First, we review opinion and evidence about the predictability of ecological systems, concluding that important, large, and exciting advances remain. We propose that these advances are constrained by lack of generally applicable and intuitive tools for assessing ecological predictability. We then introduce such a tool: the ecological forecast horizon, and suggest it as a hub for research about ecological predictability, as well as a tool for intuitively communicating the same. We provide case studies of how various sources of uncertainty and organismal characteristics influence forecast horizons. We then provide a road map for advancing ecological predictability research via ecological forecast horizons and more generally.

### 2.1 Existing knowledge about ecological predictability

Recent reviews and commentaries are optimistic about the possibility of making useful ecological forecasts (Sutherland 2006; Purves & Pacala 2008; Evans *et al.* 2013; Purves *et al.* 2013). Advances in data collection and handling, coupled with advanced quantitative methods, will enable models that provide useful predictions. Forecasts of influenza dynamics support this standpoint: despite the non-linearity and intrinsically chaotic nature of infectious disease dynamics, the timing of a disease outbreak peak was predicted up to seven weeks in advance (Shaman & Karspeck 2012). Models of population (e.g. Brook *et al.* 2000), community (e.g. Wollrab *et al.* 2012; Hudson & Reuman 2013), and ecosystem (e.g. Harfoot *et al.* 2014; Seferian *et al.* 2014) dynamics also demonstrate the predictive potential of process-based models, including individual based models (Stillman *et al.* 2015). Timely assessment of ecosystem states (Asner 2009; Loarie *et al.* 2009) and advances in hind-, now-, and forecasting methods (Dobrowski & Thorne 2011; Stigall 2012) have even allowed process-based models of land-atmosphere interactions.

Less optimistic viewpoints exist. Beckage *et al.* (2011) argue that ecological systems have low intrinsic predictability because a species’ niche is difficult to specify, because ecological systems are complex, and because novel system states can be created (e.g. by ecological engineering). Coreau *et al.* (2009) give a somewhat similar list of difficulties. These features make ecological systems ‘computationally irreducible’, such that there is no substitute for observing the real thing. Furthermore, evolution may be an intrinsically chaotic process, thus limiting the long-term predictability of ecological systems (Doebeli & Ispolatov 2014). If so, ecological responses to anthropogenic climate change are likely to be intrinsically unpredictable.

The theoretical discovery of chaos led to pessimism about forecasting. Even completely deterministic systems could have very limited forecast horizons due to the pathological sensitivity of dynamics to initial conditions. The population dynamics of a laboratory-based aquatic community were predictable only to 15–30 days due to chaotic dynamics, implying “that the long-term prediction of species abundances can be fundamentally impossible” (Benincà *et al.* 2008). Chaos also magnifies non-modelled processes (e.g. stochasticity) (Ellner & Turchin 1995), and is more common in higher dimensional systems such as ecological systems (Turchin 2003).

Other evidence about predictability comes from theoretical and empirical studies about interspecific effects. For instance, Yodzis (1988) studied whether the effects of press perturbations were directionally determined. He defined a prediction (e.g. algal biomass increases due to the addition of fish) as being directionally determined when its sign was consistent in at least 95% of cases. Yodzis found that the effects of press perturbations were frequently directionally undetermined, due to uncertainty in the parameter values. Yodzis’ findings paint a depressing picture of predicting ecological dynamics. Uncertainty in parameter values (e.g. interaction strengths) interacts with complexity (which creates indirect effects), making “implementing conservation and management strategies difficult because the effects of a species loss or an environmental perturbation become difficult to predict a priori” (quote from Wootton 2002).

Recent extensions and explanations of Yodzis’ findings provide reasons for optimism and pessimism (Novak *et al.* 2011). First, some effects of press perturbations are determined (Dambacher *et al.* 2002; Aufderheide *et al.* 2013), though these reduce in number with increases in ecological complexity (species richness and connectance of a food web) (Dambacher *et al.* 2003; Novak *et al.* 2011). Some empirical studies suggest complexity begets predictability (McGrady-Steed & Harris 1997; Berlow *et al.* 2009) while others do not (France & Duffy 2006). Second, it seems that interaction strengths can be estimated with sufficient accuracy to provide determinacy, although the demands on accuracy increase as the complexity of the ecological system increases (Novak *et al.* 2011; Carrara *et al.* 2015). Third, the results of some experimental studies have been well predicted (Vandermeer 1969; Wootton 2002, 2004). Fourth, little is know about the predictability of ecological dynamics in changing environments, such that great advances remain to be made. Fifth, predictions at the community and ecosystem level may still be possible, even if predictions at population level are not.

Whether these results and views are contradictory is unclear. Reductions in uncertainty will increase predictability, but little is known about how computationally irreducible real ecological communities are, whether different state variables (e.g. population size versus ecosystem processes) have different predictability, or about the predictability of effects of different types of environmental change (though see Fussmann *et al.* 2014; Gilbert *et al.* 2014). Ecologists must systematically and thoroughly address these challenges (Clark *et al.* 2001), though they might lack the tools needed to do so. We believe that a standard, flexible, quantitative, intuitive, and policy-relevant method for assessing ecological predictability, such as the ecological forecast horizon, will greatly aid research and communication.

### 2.2 The ecological forecast horizon

The prediction / forecast horizon as a concept goes back at least to Lorenz (1965), who wrote about how the ability to predict the weather is related to “the amount of time in advance for which the prediction is made”. Thus a forecast horizon is how far into the future (or dimensions other than time, e.g. space, phylogeny, environment) sufficiently good predictions can be made. A common reflection of the forecast horizon concept is the observation that weather forecasts are usually only made up to a specific time period into the future. After that, predictions are not good enough to be useful. However, the notion of a dynamically changing forecast horizon is important: over the past decades, the forecast horizon of ‘weather’ has increased via external effects (e.g. increase in computational power) as well as by internally optimizing the forecast system (e.g. ensemble forecasting, data assimilation, Kalman filtering).

Quantifying a forecast horizon requires a measure of how good a forecast is (we term this the *forecast proficiency*) and a *forecast proficiency threshold* above which predictions are good enough, and below which forecasts are not good enough (below we deal with how the threshold can be set). The forecast horizon is the time at which average forecast proficiency drops below this threshold (figure 1). A far forecast horizon indicates greater ability to predict (high realised predictability), a close one a weaker ability to predict (low realised predictability).

**Figure 1.**
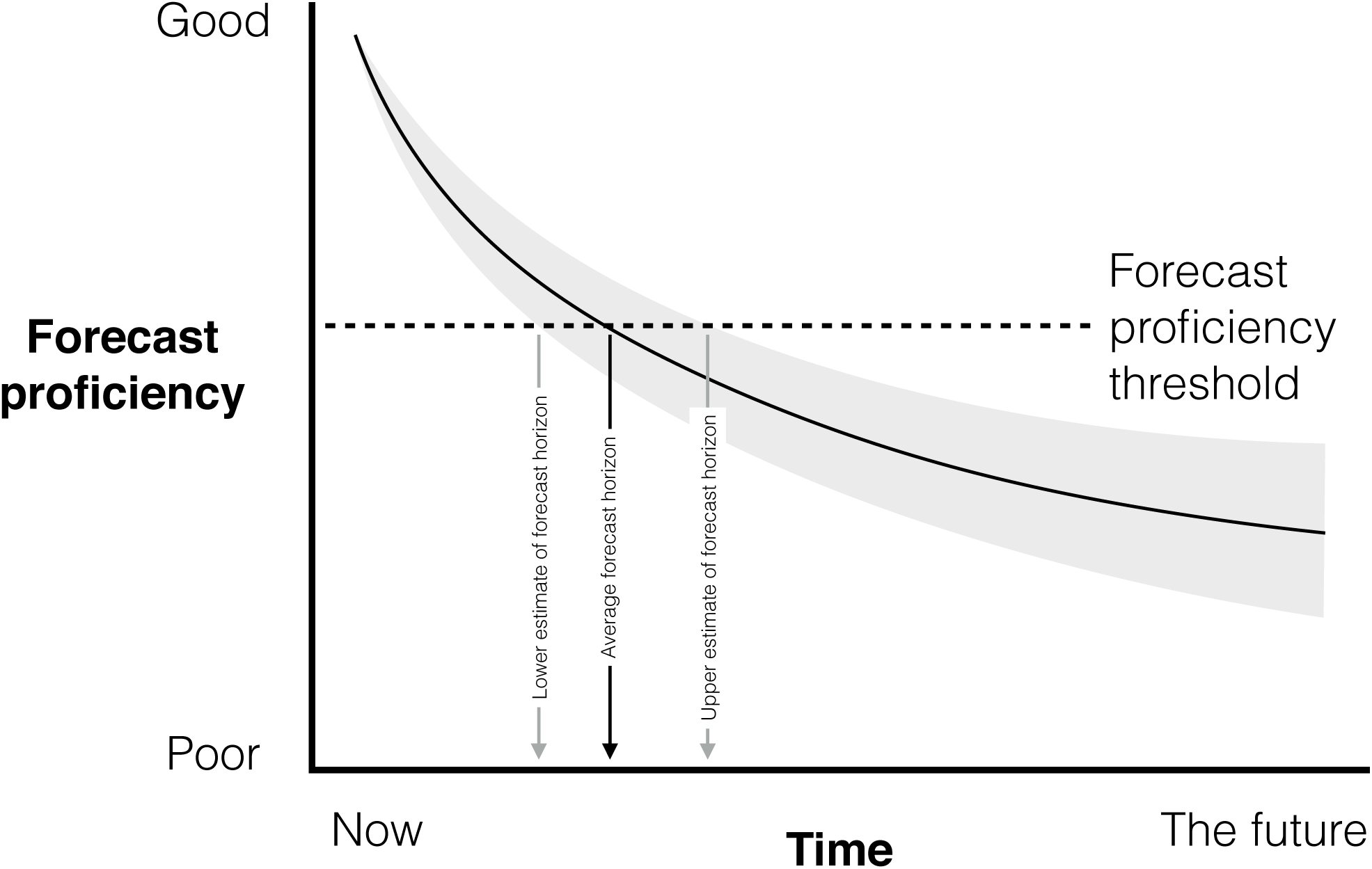
The forecast horizon is the time at which average forecast proficiency (black curved line) falls below the prediction proficiency threshold. Because forecast proficiency at any particular time will be a distribution (e.g. grey area) there will be a distribution of forecasts horizons that can be used as an estimate of uncertainty in forecast horizon (e.g. give a lower and upper estimate of the forecast horizon).

It is important to stress that there will usually be multiple possible forecasts (e.g. given uncertainty in parameter estimates or if the model contains some stochastic processes), each with a particular forecast proficiency. This will result in a distribution of forecast proficiencies and horizons (figure 1). Integrating information about these distributions into analyses and communications is important and, at least in the following case studies, is relatively straightforward.

## 3 Case studies

We provide five case studies. Two involve analyses of models, three of empirical data. Three studies involve temporal forecast horizons (how far into the future can useful forecasts be made), one spatial forecast horizon (how far away in space can useful forecasts be made), and one phylogenetic forecast horizon (how far across a phylogeny can useful forecasts be made). The temporal case studies include analyses of a simple model, a more complex model, and a complex empirical food web, and illustrate how various sources of uncertainty can impact forecast horizons. The five studies include process-based and statistical predictive models.

### 3.1 Chaos and demographic stochasticity

Using a model, we can produce a time series that we can assume is the truth. We can also produce a time series that we can assume is a forecast. If the model used to make the forecast is different from the one used to make the truth (e.g. in initial conditions, structure or parameter values), the true time series and the forecast time series can differ. This difference between time series is the forecast proficiency of the predictive model, and could be any of many quantitative measures of difference (see later). Here, we use the correlation coefficient for a window of the time series. Moving this window provides measures of forecast proficiency as a function of how far into the future the forecast is made. Note that a fully deterministic model with no uncertainty in parameter values or initial conditions will result in no difference between the truth and the prediction (i.e. an infinite forecast horizon).

We illustrate this approach with the Ricker model in the chaotic dynamic regime, as this is a simple model that can produce non-trivial behaviour. We examined the effects on forecast horizons of uncertainty in the following values: the intrinsic growth rate (*r*), the initial population size (*N*_0_), and the rate of change in carrying capacity (*K_step*). We also examined the effects of the presence or absence of demographic stochasticity in the model used to make the true time series. For each level of uncertainty in *r*, *N*_0_, and *K_step*, we drew a random value of *r*, *N*_0_, *K_step*, simulated dynamics, and calculated the forecast proficiency and forecast horizon of population dynamics. We then calculated average forecast proficiency and the average of the forecast horizon across simulations. The simulation code is available at: https://github.com/opetchey/ecopredtools.

The forecast proficiency started high (the correlation between true and predicted population size was close to 1), and dropped to near zero by at most 30 generations (figure 2). This is consistent with the chaotic nature of the model (see Box 1). Higher uncertainty in the growth rate *r*, initial population *N*_0_, or rate of environmental change *K_step* resulted in an earlier drop in forecast proficiency, compared to when there was low uncertainty. The presence of demographic stochasticity caused early and precipitous drops in forecast proficiency.

**Figure 2.**
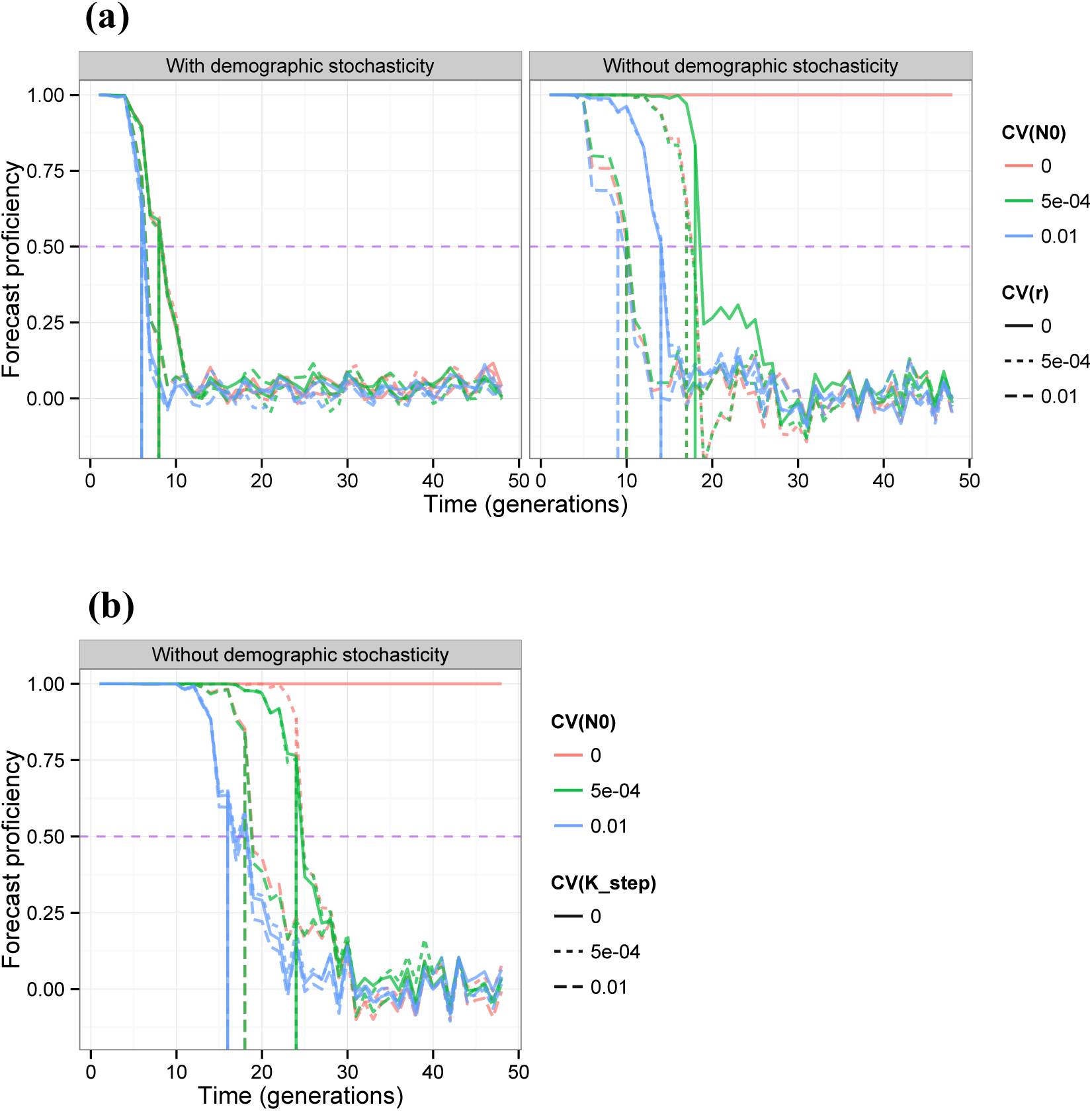
(a) Forecast proficiency as a function of how far into the future forecasts are made, for different levels of uncertainty in the growth rate parameter [CV(*r*)] of the predictive model, and uncertainty in the initial population size [CV(*N*_0_)] of the predictive model. Also shown is the effect of the presence or absence of demographic stochasticity in the true dynamics. The y-axis shows average forecast proficiencies across replicates. The horizontal purple dashed line is the forecast proficiency threshold (arbitrarily 0.3) and the vertical lines show the furthest time into the future at which forecast proficiency is above the forecast proficiency threshold, i.e. vertical lines show the forecast horizon. (b) The effect of uncertainty in the rate of environmental change (CV(K_step) relative to uncertainty in initial conditions, in the absence of demographic stochasticity.

##### 6 Text Box 1 Glossary

Accuracy: The difference between an observed and a predicted value. High accuracy implies good prediction and low accuracy poor prediction. Accuracy is an important component of forecast proficiency (see below).
Precision: The amount of uncertainty in predictions. Precise predictions will have low uncertainty (i.e. be closely grouped around the mean prediction). Imprecise predictions will have high uncertainty. Unlike accuracy, very high precision may indicate a poor predictive model that might result, for example, from failing to include a stochastic process. Low precision is also a sign of a poor predictive model. Hence, it is best if a predictive model produces a prediction that has the same uncertainty as the real system being modelled.
Uncertainty: Regan *et al.* (2002) give two classes of uncertainty: epistemic and linguistic. Epistemic uncertainty is lack of knowledge in the state of a system, for example in parameter values, processes operating, representation of processes, system components, and inherent randomness (also see Clark *et al.* 2001). See Gregr & Chan (Gregr & Chan 2014) for a discussion of the relationship between modelling assumptions and uncertainties.
Intrinsic and realised predictability: Beckage *et al.* (2011) recognise two types of predictability: the intrinsic predictability of a system, and the realised predictability achieved by a particular model of the system. The intrinsic predictability of a system is the predictability of the best possible model of that system, i.e. it is the greatest achievable predictability. Low realised predictability and high intrinsic predictability implies problems with the predictive model, such as uncertainty in parameter values. High predictability requires an intrinsically predictable system, and low uncertainty about the processes governing the system. A fully deterministic system has perfect intrinsic predictability, since perfect knowledge of parameters and initial conditions results in perfect predictions. A fully deterministic system may, however, be computationally irreducible.
Forecast proficiency: A measure of how useful a forecast is, usually some function of accuracy and or precision. We first thought to use instead the term *forecast skill*, which comes from meteorology and there usually refers to a specific measure of accuracy, mean square error, and has already been used in environmental science to assess forecasts of marine net primary production (Seferian *et al.* 2014). Forecast skill is, however, often used to mean one measure, mean square error, and we do not wish to be so specific. We propose that in ecology, the term *forecast proficiency* be general, such that any measure of accuracy or match in precision can be a measure of forecast proficiency. Thus, a model with high accuracy and appropriate precision will have high forecast proficiency. Very high precision or very low precision may both be inappropriate and contribute to lower forecast proficiency. (See Section 4.1 for a brief discussion of specific measures of forecast proficiency).
Forecast horizon: The distance in time, space, or environmental parameters at which forecast proficiency falls below the forecast proficiency threshold. Forecast horizon is closely related to concepts such as mean and maximal forecast time (e.g. Salvino *et al.* 1995).
Forecast proficiency threshold: The value of forecast proficiency above which forecasts are useful, and below which forecasts are not useful.
Retrodiction / postdiction / hindcasting: Each relates to the practice of testing the predictions of models / theories against observations already in existence at the time when the predictions were made. While care is required to understand how the existing observation might have influenced the predictions, prediction horizons can be calculated, and provide an indication about prediction into the future.

Effects of uncertainty in *r* and *N*_0_ interact (figure 3). For example, high uncertainty in *r* results in close forecast horizons regardless of uncertainty in *N*_0_, while lower uncertainty in *r* allows lower uncertainty in *N*_0_ to give farther forecast horizons. Demographic stochasticity in the true dynamics gave a very close forecast horizon, regardless of other uncertainties.

**Figure 3.**
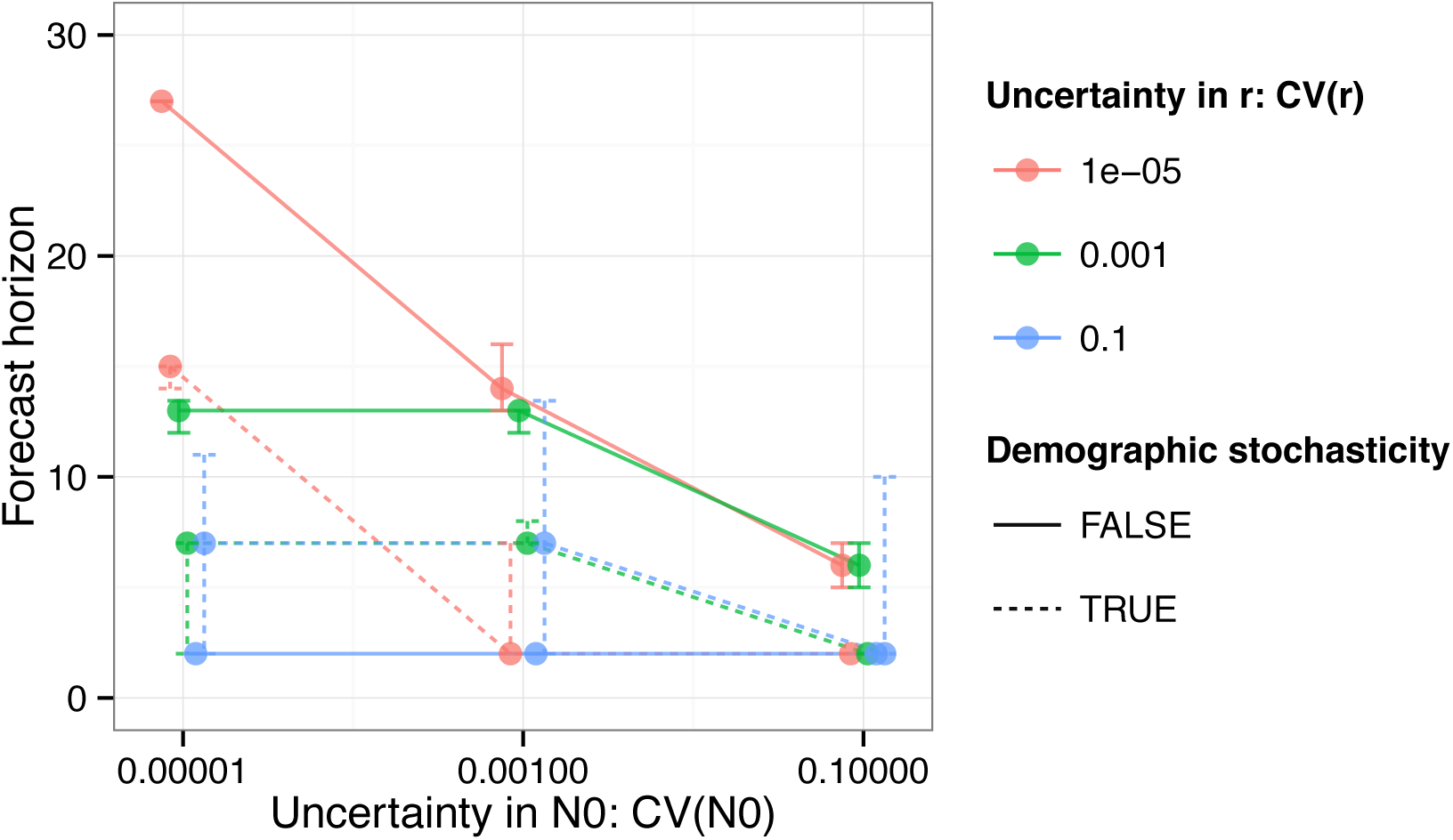
Median (+- 55^th^ to 65^th^ percentile) forecast horizon (number of generations) as a function of uncertainty in initial condition *N*0 and growth rate *r* for population dynamics with or without demographic stochasticity. The 55^th^ to 66^th^ percentile was chosen to give reasonably small error bars, for clarity.

### 3.2 Level of organisation, evolution, and environmental uncertainty

We applied the same general approach to a model of a competitive community which included evolutionary change, similar to that in Ripa *et al.* (2009). Briefly, each competing species had a trait value that determined its resource use requirements. Ecological dynamics resulted from resource depletion and therefore competition among the species, while evolutionary dynamics resulted from changes in trait values of a species (e.g. body size and resource uptake characteristics). The model also included environmental variability, implemented as random variation in the resource distribution. We evaluated the forecast proficiency of two variables, the abundance of one of the species and the total biomass of all species. We manipulated whether evolution operated in the model used to produce the true data, and also the amount of uncertainty about the nature of the environmental variability (which resulted both from intrinsic stochasticity in environmental conditions and imperfect knowledge of these conditions). Evolution was never included in the model used to forecast.

In the absence of evolution, forecast horizons for species abundance and total community biomass were very similar (figure 4). In the presence of evolution, forecast horizons were consistently farther for total community biomass. This may result from density compensation among the competing species, enhanced by supply of diversity by evolution, creating more predictable dynamics of total community biomass (e.g. Yachi & Loreau 1999). Unsurprisingly, forecast horizons are closer when there is greater uncertainty about future environmental conditions. Subsequent studies could examine the relative importance of different sources of uncertainty about environmental variability.

**Figure 4.**
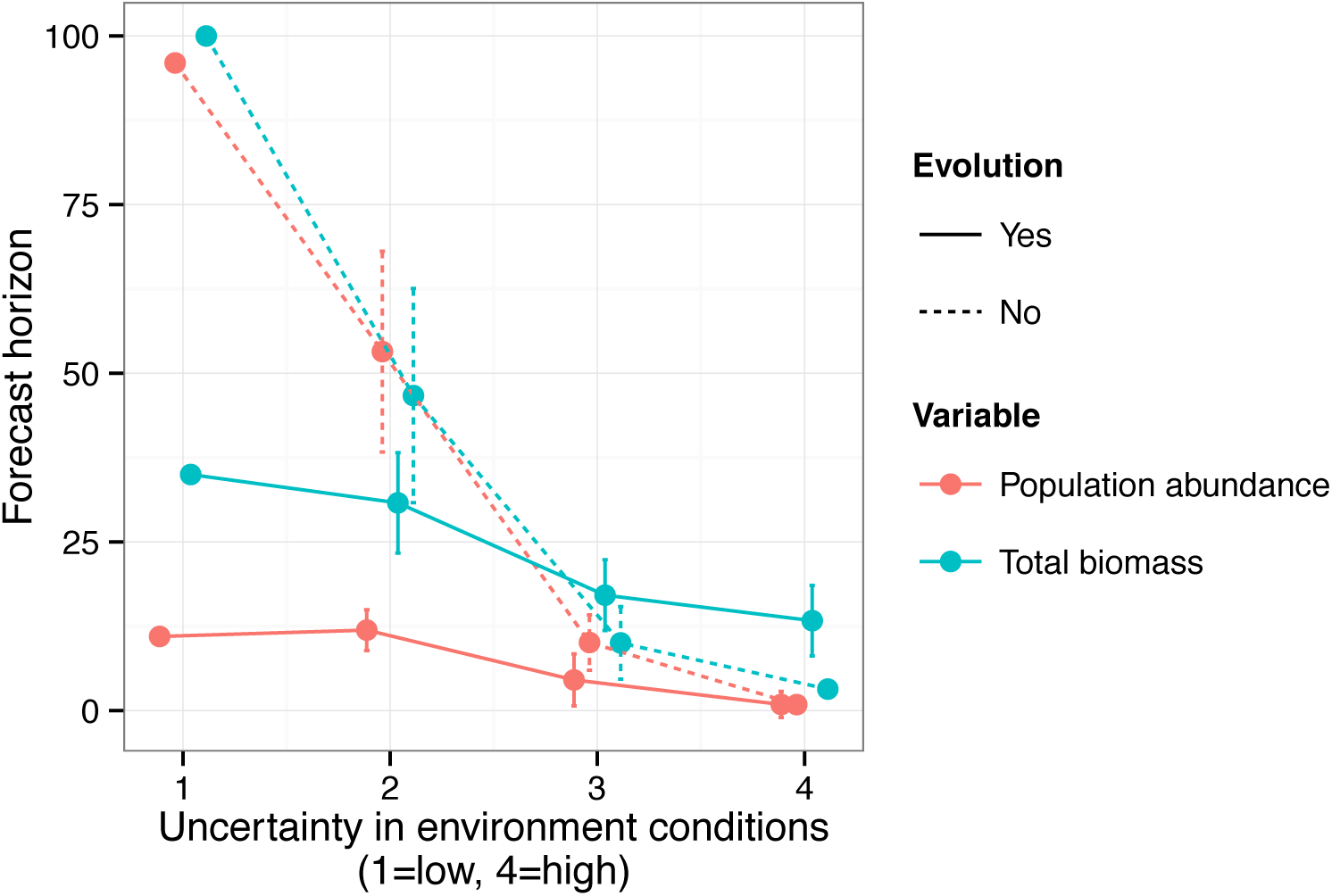
Effects of uncertainty about future environment (x-axis), of evolution, and of level of ecological organisation on forecast horizon (number of generations). Data come from a simulation study of a community of competitors. Error bars are one standard deviation. There are no error bars when there is no uncertainty in environmental conditions as then the prediction model uses the same series of environmental conditions among replicate simulations (and these are exactly the same series as used to create the “true” dynamics). Some of the error bars at high levels of uncertainty are too small to view.

### 3.3 Dynamics of an aquatic food web

A phytoplankton community isolated from the Baltic Sea was kept in a laboratory mesocosm for about eight years. Nutrients and the abundance of organisms in ten functional groups were sampled 690 times (Benincà *et al.* 2008). This long ecological time series exhibited characteristics consistent with chaos. A neural network model (correlative [statistical] rather than process-based) of the community displayed high predictability (0.70 to 0.90; measured as r-squared between observed and predicted data) in the short term only.

We extended the published study by examining variation in ecological forecast horizons among the ten functional groups and two nutrients. Forecast horizons were calculated by fitting a curve to the forecast proficiency (measured by r–squared)–forecast time relationships in Figure 2 of Benincà *et al.* (2008), and estimating the time at which forecast proficiency dropped below an arbitrarily determined forecast proficiency threshold of 0.6. Body size ranges represented by organisms in each taxonomic group were gathered from literature and online sources.

Forecast horizons exhibited a triangular relationship with organism size, with only low forecast horizons for smaller organisms and a wide range of forecast horizons for larger organisms (Figure 5a). The forecast horizon was somewhat shorter for taxa with a greater number of trophic links to other organisms (Figure 5b). The lowest p-value we were able to generate was 0.055 for the relationship between forecast horizon and number of trophic links (this value was 0.09 using size estimates provided by R. Heerkloss.) The analysis code is available at https://github.com/opetchey/ecopredtools.

**Figure 5.**
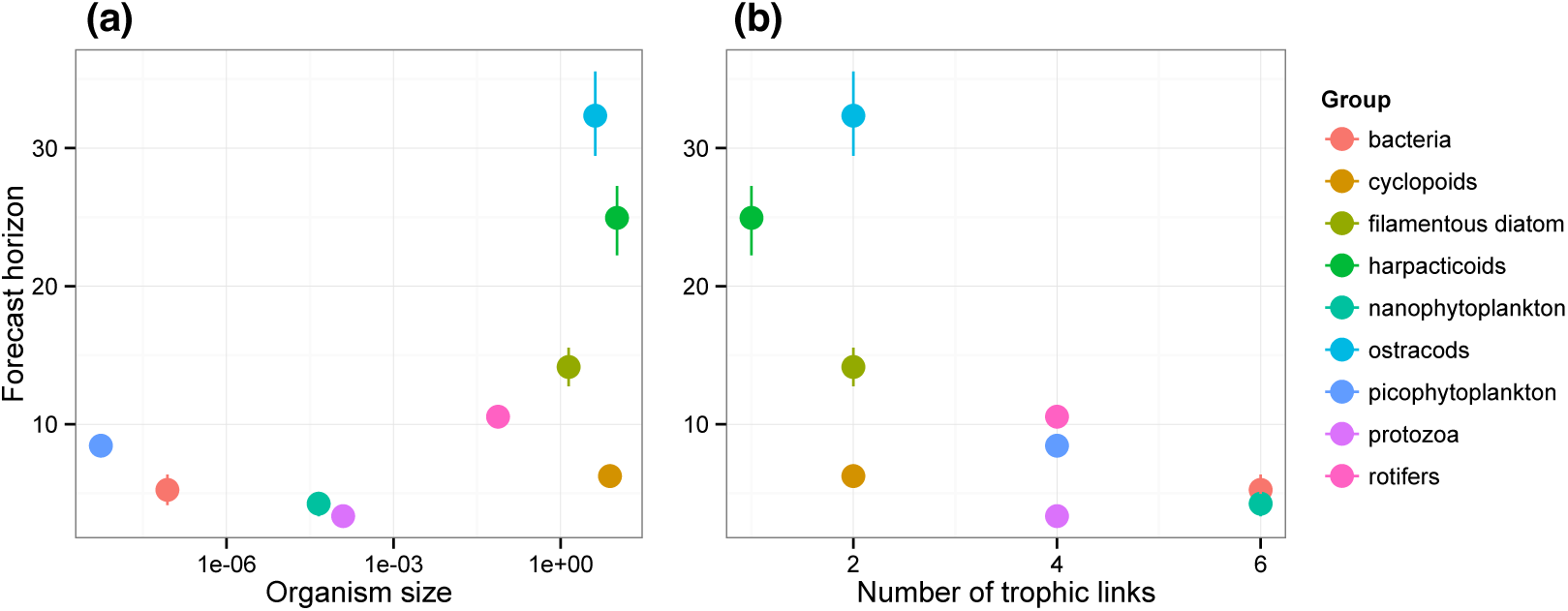
Forecast horizons (days) from Benincà *et al.* (2008) plotted against (a) approximate body size of the organisms in taxonomic groups (gathered from the literature) and (b) number of trophic links (taken from figure 1a of Benincà *et al.* (2008)). Y-error bars show the range of forecast horizons constructed from the 95% confidence intervals of curve fits to data in Figure 2 of Benincà *et al.* (2008).

Generally, longer generation times of larger organisms may partially explain this (albeit non-significant) result, though their generally smaller population sizes should increase the importance of demographic stochasticity, making for nearer forecast horizons. Hence, we do not feel confident, based on verbal arguments, about making a hypothesis regarding the expected relationship between body size and forecast horizon. The trend towards nearer forecast horizons for organisms with a greater number of trophic links may reflect the negative effects of complexity on predictability (Dambacher *et al.* 2003; Novak *et al.* 2011), perhaps related to processes linking complexity and stability (e.g. McCann 2000; May 2001).

### 3.4 Spatial forecast horizons

Forecast horizons can be made in space (maximum distance predicted to acceptable proficiency) and when the predictive model is statistical rather than process-based. A well known macroecological pattern, the decay of compositional similarity with distance curve (Nekola & White 1999; Nekola & McGill 2014), provides an example. A decay of similarity curve shows some measure of community similarity between pairs of communities on the y-axis plotted against the geographical distance between the communities (figure 6a). Sørensen similarity provides a measure of the percentage of correctly predicted species occurrences. Thus the curve provides the expected or average similarity (which can also be treated as a measure of forecasting efficiency giving the % of species correctly predicted in a community) as a function of distance. The spatial forecast horizon is the geographical distance beyond which prediction proficiency falls below a threshold (figure 6a), and in this specific example, the forecast horizon is 600km (with a threshold forecast proficiency of 0.7 correlation). Spatial forecast horizons could readily be applied to species distribution models (e.g. Pottier *et al.* 2014).

**Figure 6.**
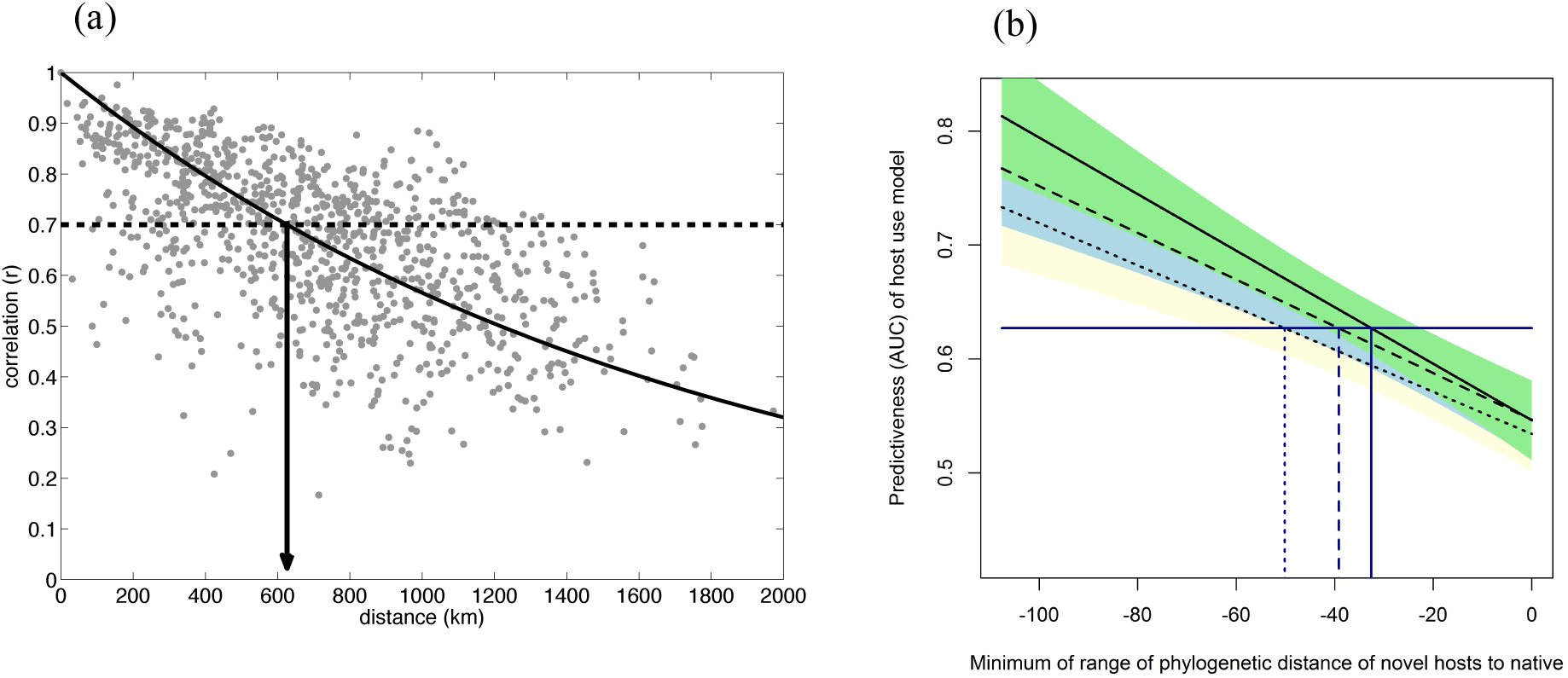
Spatial and phylogenetic forecast horizons. (a) Distance-decay of similarity in community composition. With a forecast proficiency threshold of 0.7 correlation, there is a forecast horizon of just over 600km. This example uses Pearson correlation of square-root transformed abundances as a measure of similarity of relative abundance between pairs of routes from the North American Breeding Bird Survey. (b) Fitted relationships between forecast proficiency (AUC) and phylogenetic distance (MYA) when all data were used to parameterise the forecasting model (solid line, green shading), when 2/3 of the data were used (dashed line, blue shading) and when 1/3 of the data were used (dotted line, yellow shading). The horizontal line is the median AUC for predictions from the full model. The prediction threshold for models built using reduced datasets occurred at a coarser phylogenetic distance, indicating that increased information allows finer predictions of host use over plant phylogeny. Fits are linear regressions and shaded areas the standard error of the regression.

### 3.5 Phylogenetic forecast horizons

Phylogenetic forecast horizons concern how far across phylogeny useful forecasts can be made. To illustrate phylogenetic forecast horizons, we analysed a previously published study of native Lepidoptera-plant interactions in Central Europe (Pearse & Altermatt 2013). We constructed a host-use model (a binomial GLM), in which the inclusion of a host plant in the diet of a herbivore was a function of the herbivore’s host breadth and the phylogenetic distance of that plant from another known host. We then used this model to predict the inclusion of plants introduced into Central Europe in the diet breadth of native herbivores. We divided predictions into 12 phylogenetic distance slices (12 was large enough to construct the forecast proficiency versus phylogenetic distance curve, but not so many to have too little data in each slice). We then calculated the area under the ROC curve (AUC, the measure of forecast proficiency) within each phylogenetic distance slice.

AUC related linearly and positively to phylogenetic distance, with higher forecast proficiency at farther phylogenetic distances (i.e. between plant families), and lower forecast proficiencies at smaller phylogenetic distances (figure 6b). Reducing the amount of data used to parameterise the forecasting model indicates that increased information allows better predictions of host use over plant phylogeny.

This phylogenetic forecast increases in predictability with increasing distance, whereas forecasts over time typically decrease in predictability with increasing time. Because many herbivorous insects consume a set of plants delimited at roughly the family-level, the forecast horizon for the prediction of a novel plant-herbivore interaction might be set at the family level, where predictions at a lower and higher taxonomic level are less inaccurate (e.g. Pearse & Altermatt 2013). Conversely, when considering the over-dispersion of plant communities, co-occurrence was unlikely among very close relatives (congeners), but this trend did not hold at higher taxonomic levels (Cavender-Bares *et al.* 2006), suggesting that the forecast horizon for co-occurrence might be at the genus-level, where predictions at higher levels of taxonomy will be inaccurate. Cleary more research is required to better document and understand phylogenetic forecast horizons.

## 4 Discussion

Although the primary purpose of the case studies is to illustrate ecological forecast horizons across a range of applications, they also provide some insights into the concept.

The first case study shows that uncertainty about parameters and initial conditions can interact (i.e. there are dependencies), such that focusing on decreasing uncertainty in single parameters may not improve forecast horizons. Knowledge about such dependencies will help plan effective strategies for increasing the distance of forecast horizons by decreasing uncertainties.

The second case study has two important findings. First, variables at different levels of ecological organisation may be more or less predictable and second, evolution, under some conditions, increases predictability. Although recent findings (e.g. Ward *et al.* 2014) may provide depressing reading about the predictability of population dynamics, one should not mistake these as saying anything about predictability at other levels of ecological organisation.

The third case study points towards benefits from research about organismal characteristics associated with predictability. Generalisations about the predictability of population dynamics will need to recognise the possible scaling relationship between predictability, organismal size, and other organismal characteristics.

The fourth and fifth case studies illustrate forecast horizons in dimensions other than time. Forecast horizons could also be used to estimate and convey predictability in environmental conditions (e.g. that species abundances can be usefully forecast for up to 5°C of warming, but not farther), ecological complexity (e.g. single species data can be employed to usefully forecast in communities with up to 6 species, but not beyond), and changes in community structure (Gotelli & Ellison 2006b). Similarly, when the traits that define an organism’s ecological niche are known, a forecast horizon may be defined along the axis of trait distance (Gravel *et al.* 2013). We have concerned ourselves so far with forecasting in single dimensions. Nevertheless, forecasts simultaneously across time, environmental conditions, ecological complexity, space, phylogeny or other dimensions are likely to be quite useful.

Cutting across the case studies is variability in the nature of the predictive model; in particular whether it is process-based (the Ricker and resource-consumer models) or statistical (a neural network, a regression, and a binomial glm). The forecast horizon provides a standard metric for comparing such differences in the predictive model, and systematic, thorough, and impartial assessments of the variability of the different models could aid our understanding of how to improve ecological predictability.

We believe these insights show only a fraction of the potential of forecast horizons in ecological research, and that they can be a general tool for assessing how well ecological variables and/or systems can be predicted. They are general in the sense that they can be applied in any situation where the value of a variable is predicted, and there is knowledge about the known or assumed true value of that variable. That is, they convert the output of any predictive model and any measure of forecast proficiency into a common currency: distance (be this distance in time, space, or environmental conditions). As such, ecological forecast horizons could be a powerful and flexible tool for answering questions about what is predictable in ecology, which methods offer the greatest predictive power, and how forecasting is changing through time (Simmons & Hollingsworth 2002). In the remainder of this article we suggest some avenues for furthering ecological predictability research.

### 4.1 A road map for ecological predictability research

Achieving better and more useful ecological predictions will likely benefit from a road map of activities (figure 8). Our roadmap has one destination, but has no single starting point, has no single path to the destination, and contains feedbacks. Such a road map does not prescribe a single and generally applicable methodological process for improving forecast horizons. Instead we provide some suggestions about individual activities and practices in this road map, and about some feedbacks. The order in which we present the activities below is approximately associated with specificity, from those focused on forecast horizons to more general ones. A complementary road map for improving predictability, focused on the terrestrial carbon cycles but with broad implications, already exists (Lou *et al.* 2014).

#### 4.1.1 Defining what a useful forecast is

Generally speaking, a useful forecast will be about an important variable and be sufficiently accurate and precise. This has at least three requirements: 1) a decision about the important variables to be predicted; 2) a measure of how closely a forecast is required to match the truth, i.e. a specific measure of forecast proficiency; and 3) a threshold forecast proficiency that defines “good enough”. We consider each in turn.

Which variables are important to predict is difficult to answer generally. Species abundances and distributions would be the answer according to one textbook definition of ecology (Begon *et al.* 1990). The sub-disciplines of ecology would have logical preferences for, for example, connectance in food web ecology (Petchey *et al.* 2010), species richness in community ecology (Algar *et al.* 2009), timing of infectious disease outbreaks in disease ecology (Shaman & Karspeck 2012), or biomass or carbon in ecosystem science (Harfoot *et al.* 2014).

It is then necessary to decide how to measure forecast proficiency. When the forecast variable is continuous, a number of calculations on the residuals ϵ_i_ (predicted minus actual or 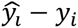) are useful, such as mean error (bias), mean square error (MSE), root mean square error (RMSE), mean absolute error (MAE), variance explained (R^2^), and correlation between predicted and observed. MSE has the useful property of combining accuracy and precision. Choices for binary variables (e.g. presence or absence, extinction or not) include the point-biserial correlation, statistics of the confusion matrix, and area under a receiver operating characteristic (ROC) curve. These vary in meaning, advantages, and disadvantages, and need to be carefully matched to purpose. For example, RMSE gives absolute error in units of the original variable while R^2^ gives relative error on a scale of 0–1 and in proportion to the total variability in the value being predicted; AUC can be misleading because the range from predicting at random to predicting perfectly is 0.5–1 (rather than the 0–1 of R^2^), which can lead people to interpret AUC scores as better than they are, and there is little intuition of what counts as a good AUC score (Bahn & McGill 2013). In situations when predicting patterns (e.g. whether dynamics are cyclic or not) is more important than exact values (Levins 1966), “pattern-oriented modelling / prediction” and associated methods for comparing predictions with data could be used (Grimm & Railsback 2012). Finally, in many predictive situations, a key issue is to ensure that the data testing the predictions are independent of the data used to calibrate the model (Bahn & McGill 2007).

Next comes a decision about the threshold forecast proficiency. For less applied research, such as that in the case studies, an arbitrary forecast proficiency threshold is sufficient, or one could use a threshold based on the average performance of a simple statistical model. Taking a more stakeholder-orientated approach, ecological forecasts and their horizons would be a service / product provided, and important variables and proficiency thresholds should be specified by stakeholders during dialogue before predictive models are employed. Such dialogues could use frameworks, including structured decision-making, to identify appropriate variables, appropriate measures of proficiency, and appropriate thresholds, given the management structure and goals of a specific problem (e.g. Guisan *et al.* 2013). Feedback between researchers and stakeholders could lead to re-evaluation of the important variables, with increased reliance on those with greater predictability.

#### 4.1.2 Complex forecast horizons

More complex situations than those in our case studies may arise. Interest in simultaneously forecasting multiple variables will require multivariate measures of forecast proficiency, perhaps aiming for one forecast horizon for all variables. Alternatively, one could calculate a forecast horizon for each variable, perhaps using variable specific-measures of forecast proficiency and forecast proficiency thresholds. The resulting set of forecast horizons could be presented individually, or combined into a single forecast horizon, depending on specific use cases.

Forecast horizons could be combined with recently developed methods for anticipating regime shifts (Scheffer *et al.* 2009). Imminent changes at the population or community level are often preceded by internal processes such as ‘critical slowing down’ in the case of population extinctions. These processes can be inferred in advance from early warning signs — in the form of generic statistical signatures — occurring after the onset of environmental perturbation and before critical system transition. The forecast horizons of such signals remain relatively unexplored.

Non-monotonic declines in forecast proficiency with forecast distance deserve further attention. They could create time windows within which useful forecasts are possible or windows in which useful forecasts are not possible (i.e. forecast blind-spots). An example of a non-monotonic relationship comes from a study of probability of quasi-extinction, in which the certainty in this probability describes a U-shape with time into the future (Ellner & Holmes 2008). This creates a prediction blind-spot, surrounded by near and far time intervals for which predictions have high certainty.

Finally, there may be situations in which it is insufficient to characterise changes in forecast proficiency with a single number (a forecast horizon). Subtleties in the relationship between forecast proficiency and time, such as when forecast proficiency is high but falls away fast versus when lower initial prediction proficiency decays slowly, are not captured by a forecast horizon (though may be in the uncertainty estimate surrounding a forecast horizon).

#### 4.1.3 Standardised tools

Providing a standardised toolbox of methods for estimating and analysing ecological predictability (including via forecast horizons) that are applicable across the diversity of ecological study and data types (e.g. experimental, observational, replicated, unreplicated) would probably be quite useful, and we are working towards developing one. Those interested in contributing should write to the corresponding author or visit our github repository (github.com/opetchey/ecopredtools).

Making connections with the numerous dynamical system theory tools that address predictability (Boffetta *et al.* 2002) is important. Box 1 shows how the forecast horizon is related to the Lyapunov exponent of a time series. Investigating the functional importance of other methods from dynamic systems theory (e.g. Salvino *et al.* 1995; Bailey 1996; Aurell *et al.* 1997; Ziehmann *et al.* 2000; Garl, *et al.* 2014) should be a research priority and will require close communication between disciplines.

#### 4.1.4 Stakeholder engagement

Harwood & Stokes (2003) proposed that ecologists face a dilemma: present persuasive simplified forecasts that pay little attention to uncertainty, or emphasise uncertainties. They go on to suggest that ecologists improve how they communicate uncertainty: “ecologists must develop rigorous methods for evaluating these uncertainties” (also see, e.g. Spiegelhalter *et al.* 2011; Raftery 2014).

Ecological forecast horizons could be an excellent tool for communicating predictability, as they are intuitive and the concept is already in common usage. One could argue they are more intuitive than other measures of predictability / uncertainty only because they hide details, such as the forecast proficiency measure. This seems to be only part of the reason, however, as one could hide details in an obscure and non-intuitive quantity. Perhaps another reason is that the quantity being communicated is a time (or distance in space, phylogeny, or environmental conditions). Furthermore, people are already familiar with the concept, for example from weather forecasting. The ease of communicating the results of quite complex research about predictability is illustrated by Shaman & Karspeck (2012) and Seferian *et al.* (2014). We emphasise, however, the need to estimate and communicate uncertainty in forecast horizons (figure 1 and vertical error bars in figures 3, 4, & 5).

Close collaboration with stakeholders is now desirable, to discover which types of stakeholders can benefit from knowing what kinds of forecast horizons. Scientific stakeholders, for example scientists that use a prediction as an input to a further model, may wish to know the forecast horizon and its consequences for predictability of their model. Scientific organisations such as IPBES (Intergovernmental Platform on Biodiversity & Ecosystem Services) may prefer to deal with forecast horizons. Other stakeholders may require other products; understanding stakeholder diversity is key to communicating uncertainty and predictability (Raftery 2014).

#### 4.1.5 Cataloguing ecological predictability

Ecologists could aim for a catalogue of important ecological variables and their ecological forecast horizons (perhaps similar to the proposal for essential biodiversity variables, Pereira *et al.* 2013). Producing this will require thorough and systematic investigations about the limits of ecological predictability. What is forecastable far into the future, what is forecastable only in the short term? Which parameters and initial conditions are more important than others in their effects on predictability? A systematic analysis of ecological forecast horizons in existing studies with appropriate data would be a worthwhile starting point to provide a baseline against which to assess improvements in ecological forecasting capabilities, as well as being useful in providing information about correlates of ecological forecast horizons (see figure 5).

Contributing to such a catalogue would require a strong commitment to model validation: “the process of determining the degree to which a model and its associated data provide an accurate representation of the real world from the perspective of the intended uses of the model” (quoted in Corley *et al.* 2014; also see Chivers *et al.* 2014). In some research fields, model verification (did we build the model correctly?) and validation (did we build the correct model?) are extremely important, and necessary for formal accreditation and use of models (for further information see Corley *et al.* 2014). Model verification and validation is relatively rare for ecological models (less than half of the disease models reported in Corley et al. (2014) experienced any model validation, a review of marine ecosystem models revealed that assumptions are mostly left implicit, and uncertainties often not considered (Gregr & Chan 2014)). Researchers and stakeholders should develop clear guidelines for verification and validation of ecological / environmental forecasting models, and decide whether accreditation is desirable.

#### 4.1.6 Improving knowledge of the governing equations

The core equations governing weather forecasting are well understood (e.g. Shuman 1978). The governing equations for ecological systems include equations linking demographic rates with environmental constraints, organismal traits and dispersal abilities, and feeding rates to resource abundances, to name only a few. The governing equations of open ecosystems (i.e. meta-ecosystems) involve movement of organisms and materials (Loreau *et al.* 2003), so that, for example, population dynamics forecast horizons are likely improved by including immigration and emigration. Optimism about the forecasting potential of process-based models rests on continued efforts to better document these and other equations governing ecological dynamics: fundamental research is necessary for improved forecasting (Courchamp *et al.* 2015). Such research should, however, be explicitly combined with research about the impacts of additional knowledge on predictive ability.

One area ripe for research is how evolution might affect ecological forecast horizons. On the one hand, incorporating the potential for evolution into simple predator-prey models might substantially increase our ability to explain ecological dynamics through time (Yoshida *et al.* 2003; Hairston *et al.* 2005; Becks *et al.* 2010; Ellner *et al.* 2011; Matthews *et al.* 2011; Fischer *et al.* 2014) and might help explore how evolution could affect transitions between different dynamic states (Ellner & Turchin 1995; Fussmann *et al.* 2000). On the other hand, evolutionary trajectories strongly influenced by ecological dynamics causing frequency-dependent selection might lead to more unpredictable evolutionary dynamics in the long term (Doebeli & Ispolatov 2014). Little is known about how such eco-evolutionary dynamics might affect the predictability of population, community, and ecosystem level responses to environmental change (but see Vincenzi 2014).

Improved knowledge about what the effects of human behaviour on predictability are, and how social systems can be coupled with ecological ones in predictive models is needed. Ecological systems include humans, such that forecasting models will need to include their actions (Palmer & Smith 2014). Scenarios coupled with quantitative models have been, and may remain, particularly important here (e.g. Cork *et al.* 2006). Furthermore, models could be used to understand the feedbacks between prediction and human intervention, whereby a prediction elicits an intervention that changes the prediction, potentially resulting in undesirable management outcomes (e.g. Peterson *et al.* 2003).

Research about the governing equations will aid our understanding of the causes of observed patterns of predictability. Are ecological systems computationally irreducible (i.e. intrinsically unpredictable) such that even the best possible parameter estimates and knowledge of initial conditions cannot provide useful forecasts? Or are ecological systems intrinsically predictable, such that feeding more and more data into models will yield continual increases in predictability?

#### 4.1.7 Statistical forecasting and autocorrelation

In the absence of sufficiently good knowledge about the governing equations, or if this knowledge is not useful for prediction (e.g. when population dynamics are chaotic), statistical models may make useful predictions. These models are representations of the autocorrelations that exist for many ecological variables in many dimensions. Autocorrelation in time and space can thus be a source of predictability, with stronger autocorrelation giving greater predictability (i.e. a farther forecast horizon). Strong autocorrelation can result in statistical models being relatively good predictors, even compared to models that contain covariates such as climate and other species (Bahn & McGill 2007). Furthermore, simple state-space reconstructions based on relatively little observed data outperform more complex mechanistic models (though see Hartig & Dormann 2013; Perretti *et al.* 2013a, 2013b) and still can distinguish causality from correlation (Sugihara *et al.* 2012). Similarly, the most accurate model of some wild animal population dynamics was the one that used the most recent observation as the forecast (Ward *et al.* 2014), and statistical models of species distributions have outperformed more mechanistic ones (Bahn & McGill 2007).

Everything else being equal, process-based models would likely be preferable, based on their suggested advantage of being able to better predict into novel conditions (Purves & Pacala 2008; Evans 2012; Schindler & Hilborn 2015). If process-based models perform less well however, then this preference may change. Statistical models at least provide a baseline of minimum forecasting proficiency, against which process-based models could be judged.

#### 4.1.8 Reducing uncertainty

Reductions in uncertainty will improve predictability alongside advances in knowledge of the governing equations. Aiming for better predictive models can even be a meeting place for these two activities, thus providing a channel by which data can inform theory and theory can inform data collection.

Careful consideration is required about whether to organise research by sources of uncertainty (e.g. parameter uncertainty, model structure uncertainties, inherent stochasticity, and uncertainty in initial condition) or by effects of ecological and evolutionary processes and variables (e.g. this paper). Particularly profitable may be a combination of both, e.g. understanding the effects of processes via their effects on uncertainties. Model validation, including sensitivity analyses, can contribute to reduce uncertainty in the parameters most important for prediction. The high predictive utility of individual based models has resulted, in part, from a focus on their validation (Stillman *et al.* 2015). Finally, parameterisation methods that use all sources of data (e.g. experiments and observations) will likely reduce uncertainties, producing more distant forecast horizons.

#### 4.1.9 Scale of predictions

Given our acknowledged poor ability to forecast environmental conditions (e.g. temperature and rainfall) even next year, ecological systems strongly controlled by environmental conditions will almost certainly show very near prediction horizons. This challenge could be overcome by predicting a moving average of system dynamics, allowing one to evaluate longer-term trends despite shorter-term uncertainty. This would be akin to predicting climate rather than weather. Research about the how predictability is related to the temporal and spatial scale of predicted variables could reveal scales of greatest (or acceptable) predictability. Such research about temporal and spatial scales would be akin to that about relationships between predictability and scale of ecological organisation (e.g. Section 3.2).

Ecological forecast horizons will likely also improve if we continue to model larger spatial extents (making the systems modelled more closed), with finer grain sizes and with more attention to modelling multiple vertical layers (e.g. below ground processes). Predictions can reasonably be expected to improve as we continue to gather data with better spatial coverage and finer resolution, and longer temporal extent data about the current and past conditions of variables of interest.

#### 4.1.10 Infrastructure improvements

Large-scale integrated investment in infrastructure for predicting ecological and ecosystem states should be considered. Ecologists, ecosystem scientists, and organisations such as IPBES should consider aiming to develop forecasting infrastructure on the scale of, for example, the UK Meteorological Office (1,800 people employed at 60 globally distributed locations, processing over 10 million weather observations a day using an advanced atmospheric model running on a high performance supercomputer, creating 3,000 tailored forecasts and briefings a day [UK Met Office web site]). Training in skills including modelling, time series analysis, working with large datasets, and communicating across traditional discipline boundaries would also be required for ecological forecasting experts.

The forecast horizon in part depends on the quality and comparability of data used to inform the predictive model. Compared to, for example, meteorology, data acquisition in the field of ecology is often less standardised across different research groups and geographic/temporal dimensions. Meteorology has used standardised tools to measure model-relevant variables, such as temperature or humidity, since the mid-19^th^ century, such that standard weather stations based on the Stevenson screen (Stevenson 1864) have been contributing comparable data across the globe for more than a century. In ecology, even basic data (e.g. following population abundances across different types of organisms) are acquired very differently across time and research groups, or are based on initiatives of individual researchers and then often lack spatial replication. Many “good” examples of time series of ecological data were actually collected without any ecological basis (e.g. records of the number of Canada lynx and snowshoe hare pelts traded by Hudson’s Bay, fisheries data, etc. which were collected mostly with an economic perspective in mind). Priority setting for which variables and parameters to measure, how to do so in a standardised way, and following explicit information standards (e.g. Darwin Core, www.tdwg.org) and ontologies may thus be of high urgency in ecology. Efforts to make such data readily accessible (Kattge *et al.* 2011; Hudson *et al.* 2014; Salguero-Gómez *et al.* 2014) in a consistent and freely available form should be redoubled (meteorological data are not only collected in a standardised way, but also made available by National Meteorological Offices) (Costello *et al.* 2013).

#### 4.1.11 Prediction competitions

Following the example of other fields with a strong interest in accurate predictions, competitions could advance methods and foster interest from non-ecologists with forecasting skills. They could provide platforms where predictions are confronted with observations on a regular basis. Being based on common datasets, they also allow direct comparisons of different methods in terms of forecasting proficiency. For instance, tests of ensembles of models (including process-based and statistical ones) compared to predictions of single methods would be possible. Such competitions are currently used in economics and are also common for improving machine learning algorithms and approaches (e.g. www.kaggle.com).

### 4.2 Conclusions

The ecological forecast horizons is a general and intuitive tool with potential to guide future research agendas to improve predictability not only by stimulating scientists to make quantitative predictions, but also by providing a mechanism to actively confront these predictions with observed dynamics. Forecast horizons provide baselines about how well we can predict specific dynamics of interest, and provide a tool for researching when and why accurate predictions succeed or fail. Given these properties, we believe that the forecast horizon can be an important tool in making the science of ecology even more predictive. Nevertheless, research should also aim for complementary, and perhaps even better, tools for advancing and organising predictability research in ecology.

## 5 Acknowledgements

Andrew Tredennick, Katie Horgan, and two anonymous reviewers provided comments that were used to improve the manuscript. Elisa Benincà provided the data from figure 2 of Benincà *et al.* (2008) and Reinhard Heerkloss provided biovolumes of taxa. Contributions of OLP, MS, and BS are supported by the University of Zurich Research Priority Program on ‘*Global Change and Biodiversity*’ (URPP GCB). Contributions of OLP are supported by SNF Project 31003A_137921 *Predicting community responses to environmental change* and two Tansley Working Groups: *FunKey Traits: Using Key Functional Traits to develop ecosystem service indicators*, and *Advancing the ecological foundations of sustainability science*. Contributions of FA are supported by the Swiss National Science Foundation Grant No PP00P3_150698. Contributions of MP are supported by the Swedish Research Council. S.K acknowledges support from the European Union’s Seventh Framework Programme (FP7/2007-2013) under grant agreement no. 283068 (CASCADE).

#### 7 Text Box 2 Lyapunov Exponents and the ecological forecast horizon

Dynamical systems theory concerns, in part, the predictability of dynamics (e.g. Boffetta *et al.* 2002). In particular, the Lyapunov exponent (LE) is closely related to intrinsic predictability of a deterministic system. The LE is a measure of the rate of separation of close trajectories (figure 7a). For example, consider the logistic map *x_t_*_+1_ = *rx_t_*(1 − *x_t_*), where *x_t_* is population size at time *t* and *r* is the growth rate. Let initial size of one replicate population be *x_o_*, and 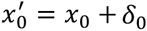 is the starting size of another population. The difference in size of the two populations initially is *δ*_0_, and the difference at time *t* is *δ_t_* (figure 7b). How *δ_t_* changes through time is characterised by the LE (*λ*), according to the equation *δ*_*t*_ = *δ*_0_*e*^*λt*^. Thus, when *λ* > 0 the intial difference grows exponentially, whereas if *λ* < 0 the difference shrinks exponentially.

**Figure 7.**
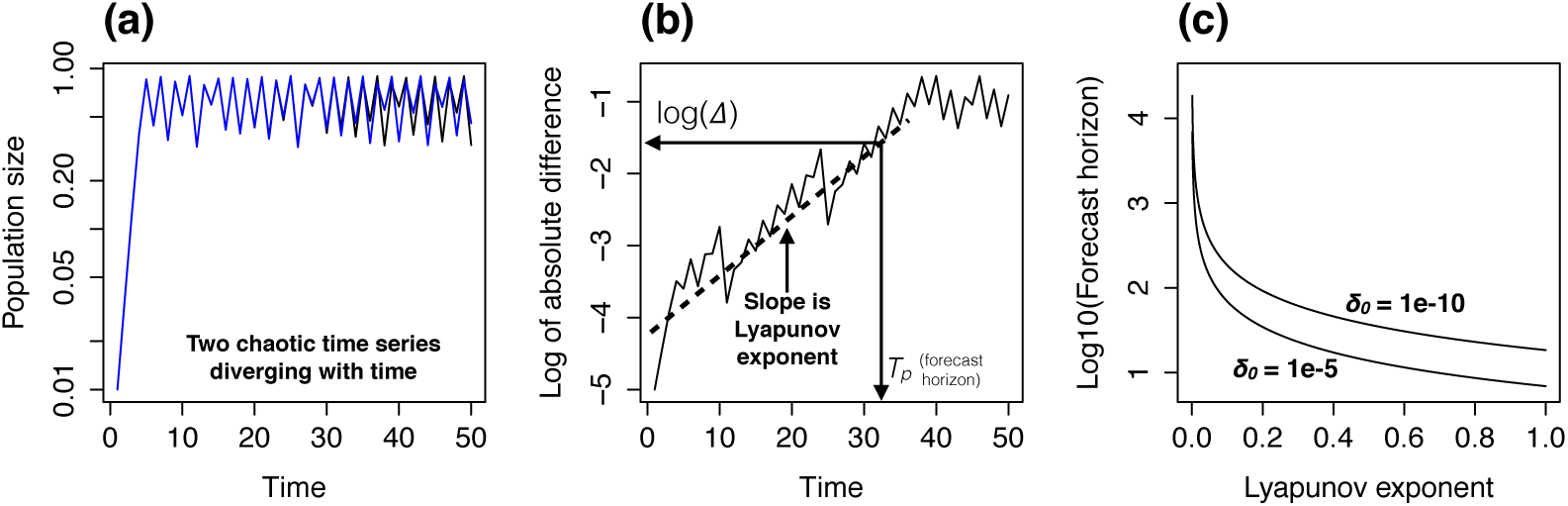
(a) Two population dynamic time series originated by two nearby initial conditions (*x*_0_ = 0.01, *x*^′^_0_ = *x*_0_ + *δ*_0_, with *δ*_0_ = 10^−5^) using the Logistic map with growth rate = 3.6. (b) Growth of the logarithm of the difference of the two times series in panel (a). (c) Relationship between forecast horizon (Tp) and the Lyapunov exponent predicted by equation 1, for two sizes of *δ*_0_.

**Figure 8.**
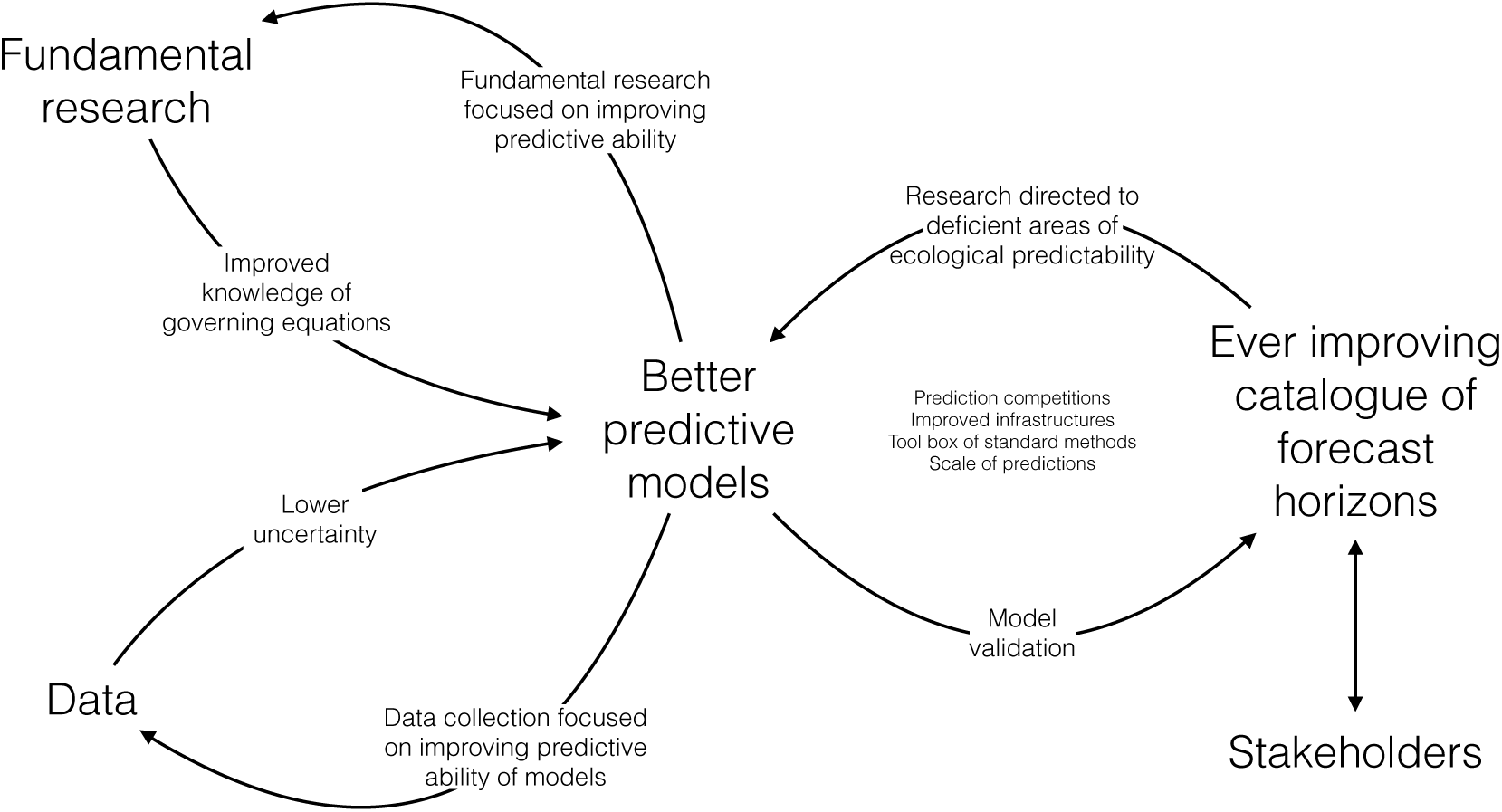
A road map for advancing ecological predictability research. Sections 4.1 and 4.2 in the main text provide details. Indirect interactions and feedbacks, such as between *Fundamental research* and *data*, are left implicit, acting via *Better predictive models*, though they are extremely important.

In order to translate the LE into a forecast horizon, we must know two things: 1) the amount of uncertainty in initial conditions (*δ*_0_); 2) the required precision of the prediction Δ (i.e. the forecast proficiency threshold). The forecast horizon is given by the heuristic equation

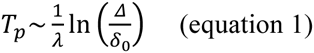

The forecast horizon *T_p_* (otherwise known as the predictability time) is the time at which a small error in the initial condition becomes large enough to preclude a useful forecast. *T_p_* is determined by the inverse of the LE, while it has weak dependence on *δ*_0_ and Δ (figure 7c). Negative LE result in an infinite forecast horizon. In case the system is multidimensional (e.g. a multispecies community) there is a LE for every dimension and predictability is determined by the largest LE of the system.

## Bibliography

1. Algar, A.C., Kharouba, H.M., Young, E.R. & Kerr, J.T. (2009). Predicting the future of species diversity: Macroecological theory, climate change, and direct tests of alternative forecasting methods. Ecography (Cop.)., 32, 22–33.

2. Asner, G.P. (2009). Tropical forest carbon assessment: Integrating satellite and airborne mapping approaches. Environ. Res. Lett., 4, 034009.

3. Aufderheide, H., Rudolf, L., Gross, T. & Lafferty, K.D. (2013). How to predict community responses to perturbations in the face of imperfect knowledge and network complexity. Proc. R. Soc. B Biol. Sci., 280, 20132355.

4. Aurell, E., Boffetta, G., Crisanti, A., Paladin, G. & Vulpiani, A. (1997). Predictability in the large: An extension of the concept of Lyapunov exponent. J. Phys. A. Math. Gen., 30, 1–26.

5. Bahn, V. & McGill, B.J. (2007). Can niche-based distribution models outperform spatial interpolation? Glob. Ecol. Biogeogr., 16, 733–742.

6. Bahn, V. & McGill, B.J. (2013). Testing the predictive performance of distribution models. Oikos, 122, 321–331.

7. Bailey, B. (1996). Local Lyapunov Exponents: Predictability depends on where you are. In: Nonlinear Dyn. Econ. Proc. Tenth Int. Symp. Econ. Theory Econom. (eds. Barnett, W.A., Kirman, A.P. & Salmon, M.). Cambridge University Press, pp. 345–360.

8. Beckage, B., Gross, L.J. & Kauffman, S. (2011). The limits to prediction in ecological systems. Ecosphere, 2, art125.

9. Becks, L., Ellner, S.P., Jones, L.E. & Hairston, N.G. (2010). Reduction of adaptive genetic diversity radically alters eco-evolutionary community dynamics. Ecol. Lett., 13, 989–997.

10. Begon, M., Harper, J.L. & Townsend, C.R. (1990). Ecology: Individuals, Populations and Communities. Blackwell Publishing.

11. Benincà, E., Huisman, J., Heerkloss, R., Jöhnk, K.D., Branco, P., Van Nes, E.H., et al. (2008). Chaos in a long-term experiment with a plankton community. Nature, 451, 822–825.

12. Berlow, E.L., Dunne, J., Martinez, N.D., Stark, P.B., Williams, R.J. & Brose, U. (2009). Simple prediction of interaction strengths in complex food webs. Proc. Natl. Acad. Sci. U. S. A., 106, 187–191.

13. Boffetta, G., Cencini, M., Falcioni, M. & Vulpiani, A. (2002). Predictability: A way to characterize complexity. Phys. Rep., 356, 367–474.

14. Brook, B.W., O’Grady, J.J., Chapman, P., Burgman, M., Akçakaya, H.R. & Frankham, R. (2000). Predictive accuracy of population viability analysis in conservation biology. Nature, 404, 385–387.

15. Cane, M.A., Eshel, G. & Buckland, R.W. (1994). Forecasting Zimbabwean maize yield using eastern equatorial Pacific sea surface temperature. Nature, 370, 204–205.

16. Cannell, M.G.R. & Smith, R.I. (1983). Thermal time, chill days and prediction of budburst in Picea Sitchensis. J. Appl. Ecol., 20, 951–963.

17. Carrara, F., Giometto, A., Seymour, M., Rinaldo, A. & Altermatt, F. (2015). Experimental evidence for strong stabilizing forces at high functional diversity of microbial communities. Ecology, in press.

18. Cavender-Bares, J., Keen, A. & Miles, B. (2006). Phylogenetic structure of floridian plant communities depends on taxonomic and spatial scale. Ecology, 87, S109–S122.

19. Chivers, C., Leung, B. & Yan, N.D. (2014). Validation and calibration of probabilistic predictions in ecology. Methods Ecol. Evol., 5, 1023–1032.

20. Clark, J.S., Carpenter, S.R., Barber, M., Collins, S., Dobson, A., Foley, J., et al. (2001). Ecological forecasts: An emerging imperative. Science (80-.)., 293, 657–660.

21. Corbet, S.A., Saville, N.M., Fussell, M., Prŷs-Jones, O.E. & Unwin, D.M. (1995). The competition box : A graphical aid to forecasting pollinator performance. J. Appl. Ecol., 32, 707–719.

22. Coreau, A., Pinay, G., Thompson, J.D., Cheptou, P.-O. & Mermet, L. (2009). The rise of research on futures in ecology: Rebalancing scenarios and predictions. Ecol. Lett., 12, 1277–1286.

23. Cork, S.J., Peterson, G.D., Bennett, E.M., Petschel-Held, G. & Zurek, M. (2006). Synthesis of the storylines. Ecol. Soc., 11, 11.

24. Corley, C.D., Pullum, L.L., Hartley, D.M., Benedum, C., Noonan, C., Rabinowitz, P.M., et al. (2014). Disease prediction models and operational readiness. PLoS One, 9, e91989.

25. Costello, M.J., Michener, W.K., Gahegan, M., Zhang, Z.-Q. & Bourne, P.E. (2013). Biodiversity data should be published, cited, and peer reviewed. Trends Ecol. Evol., 28, 454–461.

26. Courchamp, F., Dunne, J.A., Le Maho, Y., May, R.M., Thébaud, C. & Hochberg, M.E. (2015). Fundamental ecology is fundamental. Trends Ecol. Evol., 30, 9–16.

27. Dambacher, J., Li, H. & Rossignol, P. (2002). Relevance of community structure in assessing indeterminacy of ecological predictions. Ecology, 83, 1372–1385.

28. Dambacher, J., Li, H. & Rossignol, P. (2003). Qualitative predictions in model ecosystems. Ecol. Modell., 161, 79–93.

29. Diez, J.M., Ibáñez, I., Miller-Rushing, A.J., Mazer, S.J., Crimmins, T.M., Crimmins, M., et al. (2012). Forecasting phenology: From species variability to community patterns. Ecol. Lett., 15, 545–553.

30. Dobrowski, S. & Thorne, J. (2011). Modeling plant ranges over 75 years of climate change in California, USA: temporal transferability and species traits. Ecol. Monogr., 81, 241–257.

31. Doebeli, M. & Ispolatov, I. (2014). Chaos and unpredictability in evolution. Evolution (N. Y)., 68, 1365–1373.

32. Ellner, S. & Turchin, P. (1995). Chaos in a noisy world: New methods and evidence from time-series analysis. Am. Nat., 145, 343–375.

33. Ellner, S.P., Geber, M. & Hairston, N.G. (2011). Does rapid evolution matter? Measuring the rate of contemporary evolution and its impacts on ecological dynamics. Ecol. Lett., 14, 603–614.

34. Ellner, S.P. & Holmes, E.E. (2008). Commentary on Holmes et al. (2007): Resolving the debate on when extinction risk is predictable. Ecol. Lett., 11, 1–5.

35. Evans, M.R. (2012). Modelling ecological systems in a changing world. Philos. Trans. R. Soc. Lond. B. Biol. Sci., 367, 181–190.

36. Evans, M.R., Bithell, M., Cornell, S.J., Dall, S., Diaz, S., Emmott, S., et al. (2013). Predictive systems ecology. Proc. R. Soc. B Biol. Sci., 280, 20131452.

37. Evans, M.R., Norris, K.J. & Benton, T.G. (2012). Predictive ecology: Systems approaches. Philos. Trans. R. Soc. Lond. B. Biol. Sci., 367, 163–169.

38. Fischer, B.B., Kwiatkowski, M., Ackermann, M., Krismer, J., Roffler, S., Suter, M.J.F., et al. (2014). Phenotypic plasticity influences the eco-evolutionary dynamics of a predator-prey system. Ecology, 95, 3080–3092.

39. France, K.E. & Duffy, J.E. (2006). Diversity and dispersal interactively affect predictability of ecosystem function. Nature, 441, 1139–1143.

40. Fussmann, G., Ellner, S., Shertzer, K. & Hairston, J.N. (2000). Crossing the Hopf bifurcation in a live predator-prey system. Science (80-.)., 290, 1358–1360.

41. Fussmann, K.E., Schwarzmüller, F., Brose, U., Jousset, A. & Rall, B.C. (2014). Ecological stability in response to warming. Nat. Clim. Chang., 4, 206–210.

42. Gao, C., Wand, H., Weng, E., Lakshmivarahan, S., Zhang, Y. & Luo, Y. (2011). Assimilation of multiple data sets with the ensemble Kalman filter to improve forecasts of forest carbon dynamics. Ecol. Appl., 21, 1461–1473.

43. Garland, J., James, R. & Bradley, E. (2014). Model-free quantification of time-series predictability. Phys. Rev. E, 90, 052910.

44. Gilbert, B., Tunney, T.D., McCann, K.S., DeLong, J.P., Vasseur, D., Savage, V., et al. (2014). A bioenergetic framework for the temperature dependence of trophic interactions. Ecol. Lett., 17, 902–914.

45. Godfray, H.C.J. & May, R.M. (2014). Open questions: Are the dynamics of ecological communities predictable? BMC Biol., 12, 22.

46. Gotelli, N.J. & Ellison, A.M. (2006a). Forecasting extinction risk with nonstationary matrix models. Ecol. Appl., 16, 51–61.

47. Gotelli, N.J. & Ellison, A.M. (2006b). Food-web models predict species abundances in response to habitat change. PLoS Biol., 4, e324.

48. Gravel, D., Poisot, T., Albouy, C., Velez, L. & Mouillot, D. (2013). Inferring food web structure from predator-prey body size relationships. Methods Ecol. Evol., 4, 1083–1090.

49. Gregr, E.J. & Chan, K.M.A. (2014). Leaps of Faith: How Implicit Assumptions Compromise the Utility of Ecosystem Models for Decision-making. Bioscience, 65, 43–54.

50. Grimm, V. & Railsback, S.F. (2012). Pattern-oriented modelling: A “multi-scope” for predictive systems ecology. Philos. Trans. R. Soc. Lond. B. Biol. Sci., 367, 298–310.

51. Guisan, A. & Thuiller, W. (2005). Predicting species distribution: Offering more than simple habitat models. Ecol. Lett., 8, 993–1009.

52. Guisan, A., Tingley, R., Baumgartner, J.B., Naujokaitis-Lewis, I., Sutcliffe, P.R., Tulloch, A.I.T., et al. (2013). Predicting species distributions for conservation decisions. Ecol. Lett., 16, 1424–1435.

53. Hairston, N.G., Ellner, S.P., Geber, M., Yoshida, T. & Fox, J. (2005). Rapid evolution and the convergence of ecological and evolutionary time. Ecol. Lett., 8, 1114–1127.

54. Hare, J.A., Alexander, M.A., Fogarty, M.J., Williams, E.H. & Scott, J.D. (2010). Forecasting the dynamics of a coastal fishery species using a coupled climate - population model. Ecol. Appl., 20, 452–464.

55. Harfoot, M.B.J., Newbold, T., Tittensor, D.P., Emmott, S., Hutton, J., Lyutsarev, V., et al. (2014). Emergent global patterns of ecosystem structure and function from a mechanistic general ecosystem model. PLoS Biol., 12, e1001841.

56. Hartig, F. & Dormann, C.F. (2013). Does model-free forecasting really outperform the true model? Proc. Natl. Acad. Sci. U. S. A., 110, E3975.

57. Harwood, J. & Stokes, K. (2003). Coping with uncertainty in ecological advice: Lessons from fisheries. Trends Ecol. Evol., 18, 617–622.

58. Homolová, L., Schaepman, M., Lamarque, P., Clevers, J.G.P.W., de Bello, F., Thuiller, W., et al. (2013). Comparison of remote sensing and plant trait-based modelling to predict ecosystem services in subalpine grasslands. Ecosphere, 5, 1–29.

59. Hudson, L. & Reuman, D. (2013). A cure for the plague of parameters: Constraining models of complex population dynamics with allometries. Proc. R. Soc. B Biol. Sci., 280, 20131901.

60. Hudson, L.N., Newbold, T., Contu, S., Hill, S.L., Lysenko, I., De Palma, A., et al. (2014). The PREDICTS database: A global database of how local terrestrial biodiversity responds to human impacts. Ecol. Evol., 4, 4701–4735.

61. Kattge, J., Díaz, S., Lavorel, S., Prentice, I.C., Leadley, P., Bönisch, G., et al. (2011). TRY - a global database of plant traits. Glob. Chang. Biol., 17, 2905–2935.

62. Komatsu, E., Fukushima, T. & Harasawa, H. (2007). A modeling approach to forecast the effect of long-term climate change on lake water quality. Ecol. Modell., 209, 351–366.

63. Kooistra, L., Wamelink, W., Schaepman-Strub, G., Schaepman, M., van Dobben, H., Aduaka, U., et al. (2008). Assessing and predicting biodiversity in a floodplain ecosystem: Assimilation of net primary production derived from imaging spectrometer data into a dynamic vegetation model. Remote Sens. Environ., 112, 2118–2130.

64. Levine, J.M. & Antonio, C.M.D. (2003). Forecasting biological invasions with international trade. Conserv. Biol., 17, 322–326.

65. Levins, R. (1966). The strategy of model building in population biology. Am. Sci., 54, 421–431.

66. Loarie, S.R., Duffy, P.B., Hamilton, H., Asner, G.P., Field, C.B. & Ackerly, D.D. (2009). The velocity of climate change. Nature, 462, 1052–1055.

67. Loreau, M., Mouquet, N. & Holt, R.D. (2003). Meta-ecosystems: a theoretical framework for a spatial ecosystem ecology. Ecol. Lett., 6, 673–679.

68. Lorenz, E. (1965). A study of the predictability of a 28-variable atmospheric model. Tellus, 17, 321–333.

69. Lou, Y., Keenan, T.F. & Smith, M.J. (2014). Predictability of the terrestrial carbon cycle. Glob. Chang. Biol.

70. Matthews, B., Narwani, A., Hausch, S., Nonaka, E., Peter, H., Yamamichi, M., et al. (2011). Toward an integration of evolutionary biology and ecosystem science. Ecol. Lett., 14, 690–701.

71. May, R.M. (2001). Stability and Complexity in Model Ecosystems. Princeton University Press.

72. McCann, K.S. (2000). The diversity-stability debate. Nature, 405, 228–233.

73. McGrady-Steed, J. & Harris, P. (1997). Biodiversity regulates ecosystem predictability. Nature, 390, 162–165.

74. Mouquet, N., Lagadeuc, Y. & Loreau, M. (2012). Ecologie prédictive and changement planétaire. Les Cah. Prospect. - Rep., 9–44.

75. Nekola, J. & White, P. (1999). The distance decay of similarity in biogeography and ecology. J. Biogeogr., 867–878.

76. Nekola, J.C. & McGill, B.J. (2014). Scale dependency in the functional form of the distance decay relationship. Ecography (Cop.)., 37, 309–320.

77. Novak, M., Wootton, J.T., Doak, D.F., Emmerson, M., Estes, J.A. & Tinker, M.T. (2011). Predicting community responses to perturbations in the face of imperfect knowledge and network complexity. Ecology, 92, 836–846.

78. Ollerenshaw, C.B. & Smith, L.P. (1969). Meteorological factors and forecasts of helminthic disease. Adv. Parasitol., 7, 283–323.

79. Palmer, P.I. & Smith, M.J. (2014). Model human adaptation to climate change. Nature, 512, 365–366.

80. Pearse, I.S. & Altermatt, F. (2013). Predicting novel trophic interactions in a non-native world. Ecol. Lett., 16, 1088–1094.

81. Pereira, H.M., Ferrier, S., Walters, M., Geller, G.N., Jongman, R.H.G., Scholes, R.J., et al. (2013). Essential biodiversity variables. Science (80-.)., 339, 277–278.

82. Perretti, C.T., Munch, S.B. & Sugihara, G. (2013a). Model-free forecasting outperforms the correct mechanistic model for simulated and experimental data. Proc. Natl. Acad. Sci. U. S. A., 110, 5253–5257.

83. Perretti, C.T., Munch, S.B. & Sugihara, G. (2013b). Reply to Hartig and Dormann : The true model myth. Proc. Natl. Acad. Sci. U. S. A., 110, 3976–3977.

84. Petchey, O., Brose, U. & Rall, B. (2010). Predicting the effects of temperature on food web connectance. Philos. Trans. R. Soc. Lond. B. Biol. Sci., 365, 2081–2091.

85. Peterson, G.D., Carpenter, S.R. & Brock, W.A. (2003). Uncertainty and the management of multistate ecosystems: An apparently rational route to collapse. Ecology, 84, 1403–1411.

86. Pottier, J., Malenovský, Z., Psomas, A., Homolová, L., Schaepman, M.E., Choler, P., et al. (2014). Modelling plant species distribution in alpine grasslands using airborne imaging spectroscopy. Biol. Lett., 10, 20140347.

87. Purves, D. & Pacala, S. (2008). Predictive models of forest dynamics. Science (80-.)., 320, 1452–1453.

88. Purves, D., Scharlemann, J., Harfoot, M., Newbold, T., Tittensor, D., Hutton, J., et al. (2013). Ecosystems: Time to model all life on earth. Nature, 493, 295–297.

89. Raftery, A.E. (2014). Use and communication of probabilistic forecasts. *arXiv*, 1408.4812.

90. Regan, H., Colyvan, M. & Burgman, M. (2002). A taxonomy and treatment of uncertainty for ecology and conservation biology. Ecol. Appl., 12, 618–628.

91. Ripa, J., Storlind, L., Lundberg, P. & Brown, J. (2009). Niche coevolution in consumer–resource communities. Evol. Ecol. Res., 11, 305–323.

92. Salguero-Gómez, R., Jones, O.R., Archer, C.R., Buckley, Y.M., Che-Castaldo, J., Caswell, H., et al. (2014). The compadre Plant Matrix Database: an open online repository for plant demography. J. Ecol., doi: 10.1111/1365–2745.12334.

93. Salvino, L.W., Cawley, R., Grebogi, C. & Yorke, J.A. (1995). Predictability of time series. Phys. Lett. A, 209, 327–332.

94. Scheffer, M., Bascompte, J., Brock, W.A., Brovkin, V., Carpenter, S.R., Dakos, V., et al. (2009). Early-warning signals for critical transitions. Nature, 461, 53–9.

95. Schimel, D.S., Asner, G.P. & Moorcroft, P. (2013). Observing changing ecological diversity in the Anthropocene. Front. Ecol. Environ., 11, 129–137.

96. Schindler, D.E. & Hilborn, R. (2015). Prediction, precaution, and policy under global change. Science (80-.)., 347, 953–954.

97. Seferian, R., Bopp, L., Gehlen, M., Swingedouw, D., Mignot, J., Guilyardi, E., et al. (2014). Multiyear predictability of tropical marine productivity. Proc. Natl. Acad. Sci. U. S. A., 111, 11646–11651.

98. Shaman, J. & Karspeck, A. (2012). Forecasting seasonal outbreaks of influenza. Proc. Natl. Acad. Sci. U. S. A., 109, 20425–20430.

99. Shuman, F. (1978). Numerical weather prediction. Bull. Am. Meteorol. Soc., 59, 5–17.

100. Simmons, A. & Hollingsworth, A. (2002). Some aspects of the improvement in skill of numerical weather prediction. Q. J. R. Meteorol. Soc., 647–677.

101. Spiegelhalter, D., Pearson, M. & Short, I. (2011). Visualizing uncertainty about the future. Science (80-.)., 333, 1393–1400.

102. Stevenson, T.C.E. (1864). New description of box for holding thermometers. J. Scottish Meteorol. Soc., 1, 122.

103. Stigall, A.L. (2012). Using ecological niche modelling to evaluate niche stability in deep time. J. Biogeogr., 39, 772–781.

104. Stillman, R.A., Railsback, S.F., Giske, J., Berger, U.T.A. & Grimm, V. (2015). Making Predictions in a Changing World: The Benefits of Individual-Based Ecology. Bioscience, 65, 140–150.

105. Sugihara, G., May, R., Ye, H., Hsieh, C., Deyle, E., Fogarty, M., et al. (2012). Detecting causality in complex ecosystems. Science (80-.)., 338, 496–500.

106. Sutherland, W.J. (2006). Predicting the ecological consequences of environmental change: A review of the methods. J. Appl. Ecol., 43, 599–616.

107. Sutherland, W.J., Armstrong-brown, S., Paul, R., Brereton, T.O.M., Brickland, J., Colin, D., et al. (2006). The identification of 100 ecological questions of high policy relevance in the UK, 617–627.

108. Tallis, H.M. & Kareiva, P. (2006). Shaping global environmental decisions using socioecological models. Trends Ecol. Evol., 21, 562–8.

109. Travis, J., Coleman, F.C., Auster, P.J., Cury, P.M., Estes, J.A., Orensanz, J., et al. (2014). Correction for Travis et al., Integrating the invisible fabric of nature into fisheries management. Proc. Natl. Acad. Sci. U. S. A., 111, 4644.

110. Turchin, P. (2003). Complex Population Dynamics: A Theoretical/Empirical Synthesis. Princeton University Press.

111. Vandermeer, J. (1969). The competitive structure of communities: An experimental approach with protozoa. Ecology, 50, 362–371.

112. Vincenzi, S. (2014). Extinction risk and eco-evolutionary dynamics in a variable environment with increasing frequency of extreme events. J. R. Soc. Interface, 11, 20140441.

113. Ward, E.J., Holmes, E.E., Thorson, J.T. & Collen, B. (2014). Complexity is costly: A meta-analysis of parametric and non-parametric methods for short-term population forecasting. Oikos, 123, 652–661.

114. Wollrab, S., Diehl, S. & De Roos, A.M. (2012). Simple rules describe bottom-up and top-down control in food webs with alternative energy pathways. Ecol. Lett., 15, 935–946.

115. Wootton, J.T. (2002). Indirect effects in complex ecosystems: Recent progress and future challenges. J. Sea Res., 48, 157–172.

116. Wootton, J.T. (2004). Markov chain models predict the consequences of experimental extinctions. Ecol. Lett., 7, 653–660.

117. Yachi, S. & Loreau, M. (1999). Biodiversity and ecosystem productivity in a fluctuating environment: The insurance hypothesis. Proc. Natl. Acad. Sci. U. S. A., 96, 1463–1468.

118. Yodzis, P. (1988). The indeterminacy of ecological interactions as perceived through perturbation experiments. Ecology, 69, 508–515.

119. Yoshida, T., Jones, L.E., Ellner, S.P., Fussmann, G.F. & Hairston, N.G. (2003). Rapid evolution drives ecological dynamics in a predator-prey system. Nature, 424, 303–306.

120. Ziehmann, C., Smith, L.A. & Kurths, J. (2000). Localized Lyapunov exponents and the prediction of predictability. Phys. Lett. A, 271, 237–251.

